# The potential of undersown species identity vs. diversity to manage disease in crops

**DOI:** 10.1101/2024.02.29.582760

**Authors:** Seraina Lisa Cappelli, Luiz Alberto Domeignoz Horta, Stephanie Gerin, Jussi Heinonsalo, Annalea Lohila, Krista Raveala, Bernhard Schmid, Rashmi Shrestha, Mikko Johannes Tiusanen, Paula Thitz, Anna-Liisa Laine

## Abstract

1. In the absence of chemical control with its negative side effects, fungal pathogens can cause large yield losses, requiring us to develop agroecosystems that are inherently disease resistant. Grassland biodiversity experiments often find plant species diversity to reduce pathogen pressure, but whether incorporating high biodiversity levels in agricultural fields have similar effects remains largely unknown.
2. We tested if undersown plant species diversity could reduce barley disease, and whether the effect was mediated through above- or belowground mechanisms, by combining an agricultural field trial with a soil transplant experiment.
3. As predicted, barley disease decreased in the presence of undersown plants. Undersown species richness had no effect, but their abundance led to early season disease reduction. Aboveground mechanisms underpinned this disease reduction. Barley yield slightly decreased with increasing undersown species richness, and undersown species varied in their impact on yield.
4. We identified two undersown species with similar functional traits that contributed most to disease reduction and had the potential to increase barley yield. Furthermore, our results indicate that aboveground mechanisms caused this. We show that agroecosystem functioning can be improved without trade-offs on yield by targeted selection of undersown species.

## Introduction

Pathogenic fungi are ubiquitous (Fisher et al., 2012) and responsible for 70% of all known plant diseases (Carris et al., 2012) causing approximately 12% of yield losses in crops, despite growing use of fungicides (Oerke, 2006; Savary et al., 2019; Walters et al., 2012). Intensive fungicide use promotes resistant pathogen evolution (Lucas et al., 2015), poses human health risks (Lushchak et al., 2018) and impacts the environment (Meena et al., 2020; Zubrod et al., 2019). Monoculture yield decline over time is often associated with the accumulation of above or belowground pathogens (Bassi et al., 2023). Alternative fungal pathogen control is needed to reduce negative impacts of fungicides and maintain their efficacy.

Intercropping, undersowing and other diversification practices are potentially powerful tools to mitigate negative impacts of intensive farming, maintain ecosystem functioning and improve food security (Brooker et al., 2015; Cappelli et al., 2022; IPBES, 2019; Kleijn et al., 2019; McAlvay et al., 2022). Species richness in agricultural fields may also reduce disease pressure in crops (Finckh et al., 2000; Karjalainen, 1986; Larkin, 2015; Trenbath, 1993), aligning with general findings of decreased disease with increasing species richness in ecological diversity experiments (X. Liu et al., 2016; Mitchell et al., 2003; Rottstock et al., 2014). In mixed cropping systems, especially cereal crops benefit from reduced disease pressure (Bedoussac et al., 2015; Zhang et al., 2019). For example, intercropping barley with different legume species decreased net blotch disease by 50% or more in a study in Denmark (Hauggaard-Nielsen et al., 2008).

Mechanisms such as decreased and intercepted pathogen dispersal, decreased host susceptibility or induced resistance reduce disease pressure in diverse plant communities (Civitello et al., 2015; Keesing et al., 2006) and contribute to reduced disease pressure in mixed cropping systems (Finckh et al., 2000; Trenbath, 1993). Disease amplification with increasing richness is possible, but less commonly observed (e.g. Ocimati et al., 2018; Power & Mitchell, 2004). Aboveground, non-host species in the plant community can physically intercept dispersing spores. For example, wheat plants have been shown to intercept spore dispersal between barley plants of a barley pathogen (Burdon & Chilvers, 1977). The canopy structure created by different plant communities can affect the microclimate in the vegetation, which can alter pathogen population dynamics. Mixing tall and short varieties of rice created a canopy that dried out more effectively, which reduced how long leaves remained wet and decreased rice blast infection (Zhu et al., 2000). Neighboring plants can also suppress pathogens through allelopathic effects on pathogen growth (Gómez-Rodríguez et al., 2003), or by triggering defense reactions in uninfected neighboring plants (Wenda-Piesik et al., 2010). With increasing plant species diversity also pathogen diversity increases while pathogen abundance decreases (Rottstock et al., 2014). Being exposed to non or low virulent microbes can trigger induced defense pathways, which can then prime the crop to be resistant against its own pathogens (Finckh et al., 2000). Exposing barley plants to two fungi that are non-pathogenic for barley reduced colonization success of net blotch causing fungus *Pyrenophora teres* by more than two thirds (Lyngs Jørgensen et al., 1998).

Similar mechanisms can also occur belowground. Soils of diverse plant communities host a higher diversity of mycorrhizal fungi, than those of species poor communities (Guzman et al., 2021) and mycorrhiza can prime crop plants to be more resistant to diseases (Pozo & Azcón-Aguilar, 2007). Volatile organic compounds of pathogen exposed barley roots slowed the growth of other barley pathogens (Fiers et al., 2013; Kaddes et al., 2016). Another belowground mechanism by which neighboring plants can affect the health of a focal species is through plant nutritional status, which can be improved in mixtures due to complementary resource use between plant species (Ashton et al., 2010, 2010; McKane et al., 2002). For example, intercropping with legumes improves cereal nutrition with nitrogen (Bedoussac et al., 2015) which can increase resistance against certain pathogens (Dordas, 2008; Tripathi et al., 2022). The soil microbial community is an important driver of soil-mediated disease suppression. Diverse plant communities harbor diverse soil microbial communities, which can promote disease resistance in plants through the production of toxins and metabolites, by competitive exclusion of pathogens, by triggering systemic defenses in the hosts, or by improving plant health through increased stress resistance (Bollmann-Giolai et al., 2022; Garbeva et al., 2004). More diverse plant communities can host larger (Eisenhauer et al., 2010), more diverse (Maciá-Vicente et al., 2023) and more beneficial soil microbial communities (Stefan et al., 2021) than monocultures.

With higher plant species richness, it is likely that more of those mechanisms occur simultaneously, and that total disease suppression increases both via above- and belowground mechanisms. Different plant species likely vary in their contribution to disease suppression in neighboring plants, as has been shown for root pathogens in grasslands (Ampt et al., 2022). Disentangling the contribution of above and belowground mechanisms to disease reduction with increasing species richness in a crop field can guide research-based agroecosystem designs.

In an agricultural context, impacts of diversification should carfully also consider effects on yield. Intercropping often increases total mixture yield compared with the average monoculture yield of the species used in the mixture (Martin-Guay et al., 2018), in the same way as it does in biodiversity experiments (Balvanera et al., 2006). However, the per-area yield of a single crop in mixture is usually lower than of the corresponding monoculture (Abu-Bakar et al., 2014; Bedoussac et al., 2015; Hauggaard-Nielsen et al., 2001). Therefore, despite all beneficial effects such as disease suppression, intercrops or undersown species can compete with the main crop and reduce its yield. Different neighbor species vary in the magnitude of this competitive impact (Abu-Bakar et al., 2014; Bedoussac et al., 2015). Especially legume intercrops have less negative impact on cereal yields than non-leguminous species (Abu-Bakar et al., 2014; Duchene et al., 2017). Similarly, undersowing with legumes reduces barley yield less than undersowing with grasses (Karlsson–Strese et al., 1998; Kunelius et al., 1992). This might be because legumes suppress other species that would compete more for soil nitrogen with a main crop, because they can access atmospheric nitrogen (Hauggaard-Nielsen et al., 2001). Intercropping has positive impacts on soil quality, including water retention, and the effects of this on yield may become apparent only in the long-term and particularly during drought episodes (Brooker et al., 2015).

In this manuscript we tested if undersown diversity reduces foliar disease in barley and if such diversity effects are driven by undersowning in general or by particular undersown species. Furthermore, we asked whether above- or belowground mechanisms were underpinning the effect of undersown diversity on foliar disease (H1 in Figure 1). We tested if undersown diversity changed the soil fungal communities towards a disease-suppressive community composition (H2 in Figure 1). Finally, we tested the impact of the undersown diversity and single undersown species on barley yield (H3 in Figure 1).

**Figure 1:**
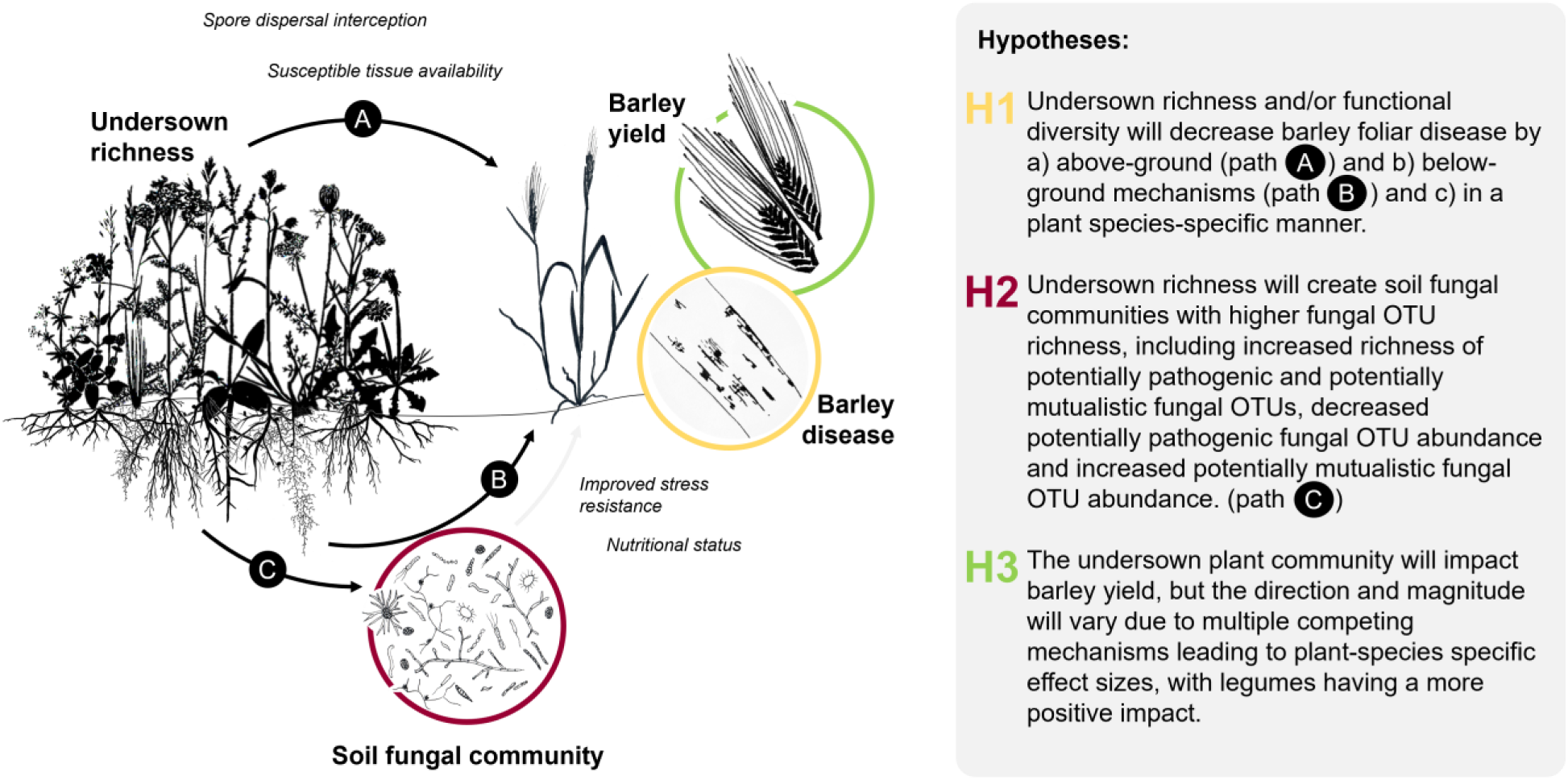
Hypotheses (H) on above- and belowground mechanisms by which plant diversity affects barley disease, yield and the composition of soil fungal communities, based on operational taxonomic units (OTU)

## Methods

### TWINWIN field experiment

The TWINWIN experiment was established in 2019 in Helsinki (60°13’N/25°01’E) to study the effects of undersown plant species and their diversity on above- and belowground agroecosystem functioning. The 30 years annual mean temperature was 5.97±0.77°C and the annual precipitation 682±115mm (Finnish Meteorological Institute, 2019). Barley (*Hordeum vulgare* L. var. Harbinger) was sown with 80kgha^-1^y^-1^ nitrogen at the end of May or beginning of June (date varied between years) in 56 out of 60 plots of 40m^2^ arranged in four blocks (Figure 2D). Four plots (not used in this study) remained without plants. Barley grew alone with (4 plots) and without herbicide (8 plots) as control (undersown species richness = 0). In the other plots, barley grew with undersown plant communities, which represented undersown richness levels one (eight single undersown species, each replicated in three plots), two (ten unique mixtures of two undersown species), four (six unique mixtures of four undersown species) and eight (one mixture of all eight undersown species, replicated in four plots) (Figure 2A). The eight undersown species differed in their ability to fix nitrogen and in their rooting depth: *Trifolium repens* L., *T. hybridum* L. (shallow rooting, nitrogen fixing), *T. pratense* L., *Medicago sativa* L. (deep rooting, nitrogen fixing), *Lolium multiflorum* Lam., *Phleum pratense* L. (shallow rooting, non-nitrogen fixing), *Festuca arundinacea* Schreb., *Cichorium intybus* L. (deep rooting, non-nitrogen fixing, Figure 2B). The species compositions in two and four species mixtures varied in functional diversity (number of functional types: ability to fix nitrogen x rooting depth) within their species richness levels (Figure 2C). Undersown plants were sown one week after barley in rows between the barley. Six weeks after barley sowing, we clipped weeds taller than barley at ground level. In early September, the barley was harvested to measure **barley yield** (H3 in Figure 1). Plots were tilled in spring before sowing for the next season.

**Figure 2:**
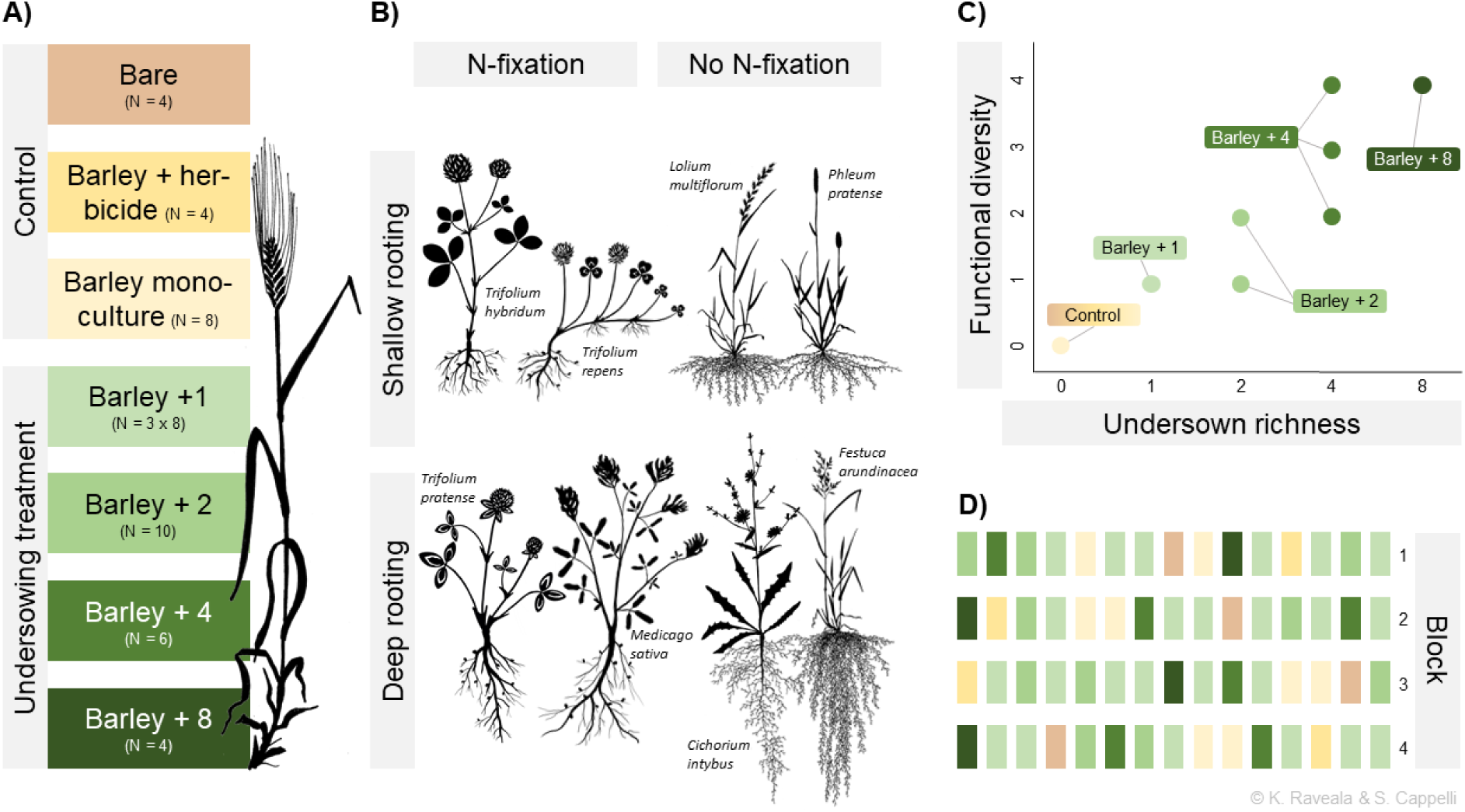
TWINWIN experiment design: A) Three control and four undersowing treatments. B) The 8 undersown species vary in rooting depth and N-fixation capacity, leading to four different “functional types”. C) The undersown communities vary in richness and functional diversity (number of functional types of undersowns in a plot). D) The 60 plots are arranged in 4 blocks (see also Table S1)

**Table 1:** replication statement.

|  | Variable of interest | Scale of inference | Scale at which the factor of interest is applied | Number of replicates at the appropriate scale |
| --- | --- | --- | --- | --- |
| TWINWIN main experiment | Disease | Leaf per individual | Plot | 3 leaves of 20 and 10 individuals per plot in July and August respectively. See Figure 2A for details on replicates per undersowing treatments. Total 56 out of 60 plots used. |
|  | Yield | Plot |  | See Figure 2A for details on replicates per undersowing treatments Total 56 out of 60 plots used. |
| Soil transplant experiment | Disease | Leaf | Pot | 3 leaves of one plant per pot, 5 replicates per environment x soil origin combination. Total 245 pots. |
|  | Soil fungal community metrics | Pot |  | 5 replicates per environment x soil origin combination. Total 231 pots. |

To verify H1 (Figure 1), we assessed barley **leaf damage** by leaf spots on the top three leaves (if available) of 20 and 10 haphazardly selected barley stems per plot in mid-July and August 2020 respectively. Per leaf, the damage was classified as 0 (0% leaf area damage), 1 (<10%), 2 (10-25%), 3 (25-50%), or 4 (50-100%) (adapted from Dirzo & Domínguez, 2010). Net blotch (*Pyrenophora teres* Drechs. 1923), a hemibiotrophic fungal pathogen of barley (Z. Liu et al., 2011) was dominant and its presence was recorded. We also visually estimated the **relative abundances** of all species in all plots at the end of July. All data used for this study was collected in 2020.

### Soil transplant experiment

To evaluate how diversity affects disease through above- and belowground mechanisms (Figure 1, H1a-b) we did a reciprocal soil transplant experiment. We grew barley plants in pots containing soil originating from different TWINWIN plots (collected in early June 2020) in the greenhouse and reciprocally placed the pots back into the different field plots (late June). We used a subset of the plots and undersown species: *L. multiflorum*, *F. arundinacea*, *T. hybridum* and *M. sativa*. We used soil from barley monoculture plots, the plots with either one of the four undersown species alone, these four undersown species together, and the plots with barley and eight undersown species, leading to seven different soil types, varying in undersown species richness (hereafter soil-origin richness). Pots with each soil type were placed back into the original plots (7 different environment types varying in undersown species richness (hereafter environment richness). We replicated each soil by environment type combination 5 times (245 pots, details in Supplementary methods).

To assess soil origin and environment richness impact on disease (Figure 1, H1a-b), we measured pathogen leaf damage on the tallest tiller per pot as described above, one and five weeks after placing the pots in the field, which is one and two months after sowing the seeds to the pots.

To test if soil-origin richness changed soil fungal communities (Figure 1, H2), we sequenced the fungal communities (ITS7 – ITS4 region) in a subset of 231 pots at the end of the experiment. Based on the FUNGuild database (Nguyen et al., 2016) we checked for each operational taxonomic unit (OTU) if it was potentially pathogenic or potentially symbiotic. Then, we calculated OTU **Shannon diversity index**, the **relative abundance of potentially pathogenic OTU reads** (contribution of pathogens to total fungal abundance), the **proportion of potentially pathogenic OTUs** (contribution of pathogens to total fungal diversity) and checked for the **presence of potentially symbiotic fungi** for each pot (Details in Supplementary methods).

### Statistical analysis

To test the hypothesis of undersown diversity leading to reduced barley disease in the TWINWIN main experiment (Figure 1, H1), we ran a series of cumulative link mixed effects models (Ordinal package, Christensen, 2019). Cumulative link mixed effects models account for the ordered categorical nature of the response variable disease, that was scored in ordered percent damage ranges (see above, Ananth & Kleinbaum, 1997). We corrected for block and leaf position (indication of leaf age) by fitting these terms before any other fixed effects and included plot and plant individual as random effects. To check if undersown richness and/or functional diversity explain disease, we fit four models with different combinations of the two metrics as fixed effects and compared AIC values of those models. For this and all following analyses the logarithm of undersown richness plus one was used, because this allowed us to include control plots with undersown richness = 0. The model including undersown richness alone had the lowest AIC value (Table S2) and therefore we continued with a model that did not include undersown functional diversity (model 1). To test if any diversity effect was driven by herbicide treatment in the plots without undersown species (zero undersown richness) or by undersowing as such, we fitted contrasts for herbicide treatments (with vs. without, model 2) or herbicide (with vs. without) and undersowing (0 vs. >0) treatments (model 3) before fitting undersown richness (testing the richness effect for plots with richness >0). To test for particular species effects (Figure 1, H1b), we used model 1 as baseline and added a presence/absence term for each undersown species (model 4). The model was simplified by stepwise removal of non-significant presence/absence terms using likelihood-ratio tests. To account for variable establishment success of the different undersown species in different plots, presence/absence data were replaced with species relative abundances and the relative abundance of weeds and barley were included as covariates in a further model (model 5). This model was also simplified.

To analyze effects of the undersown treatments on barley yield in the TWINWIN main experiment (H3), yield was analyzed in response to undersown richness and/or functional diversity, again controlling for block (model 6), using linear models. Again, we used AIC comparisons to find the best combination of undersown diversity metrics and the model with undersown richness alone had the lowest AIC value (Table S11) and was used for detailed analysis (model 1). Then, we fitted two additional models in which we first corrected for herbicide treatment (model 7) or for herbicide and undersowing treatments (model 8). Furthermore, we checked if disease reduction by undersown richness explained a potential diversity effect, by fitting the proportion of leaves with high disease (>10% leaf area damaged) as covariate before fitting undersown richness, which was not the case (data not shown).

To test the effects of single undersown species on yield (H3c), we fit another linear model with a treatment variable that distinguishes between plots in which barley was grown alone, alone with herbicide, together with any of the undersown species (undersown diversity of 1 per species), and with two, four and eight undersown species. Again, block was fit before treatment (model 9). We analyzed pairwise differences between treatments with Tukey’s HSD post-hoc test (emmeans package, Lenth, 2022).

Disease in the soil transplant experiment was analyzed per time point using cumulative link mixed effects models (ordinal package, Christensen, 2019) with leaf position, soil-origin richness, environment richness and the interaction between the two as fixed effects and pot as random effect (model 10, H1a-b). For this and all following analyses the logarithm of environment and soil-origin richness plus one were used.

To test if the treatments affected the soil fungal community composition in the soil transplant experiment we analyzed all metrics in response to environment and soil-origin richness and their interaction, including plot as random effect (H2). OTU Shannon diversity was analyzed with a linear mixed effects model (model 11, lme4 package, Bates et al., 2015). The proportion of potentially pathogenic OTUs and the relative abundance of potentially pathogenic OTU reads were analyzed with generalized linear mixed models with beta-binomial distribution (model 12 and 13, glmmTMB package, Brooks et al., 2017). We analyzed presence/absence of potentially symbiotic OTUs with a generalized linear mixed effects model with a binomial distribution (model 14, lme4 package, Bates et al., 2015). We also used cumulative link models to test if the fungal community metrics explained disease levels, but none of them had significant effects (results not shown). All analyses were done in R (Version 4.0.2, R Core Team, 2022).

## Results

### TWINWIN field experiment

Ninety-six percent of the 3312 leaves and 100% of the 1120 surveyed barley individuals in the TWINWIN experiment showed some signs of **disease** when we assessed disease in July 2020. We found distinct symptoms of net blotch disease caused by *Pyrenophora teres* on 44% of the individuals. Likely, its proportion was higher, but we could not with certainty identify the damage as net blotch when the lesions were small. It is likely that pathogens causing other leaf spot diseases like *Cochliobolus sativus* were present, too. In August 2020, all assessed leaves (total 1666) showed signs of infection and most (93%) were in the highest damage category, which made detailed analysis impossible. The following results are based on the disease measurements in July.

Undersown richness explained barley foliar disease better than undersown functional diversity (Table S2) and we therefore did not include functional diversity in the final models. After correcting for block and leaf position, undersown richness had a marginally significant negative effect on disease (p=0.064, Table S3-S4, Figure S2, model 1). This effect was largely explained by the difference between plots with and without undersown species (Table S3-S4, Figure S3c, model 3). Undersowing reduced the probability of heavy damaged (>10% of the leaf area damaged) from 70.38% to 18.95% (p=0.007, Table S3-S4, Figure S3b, model 3, H1). When including the presence of single undersown species instead of herbicide and undersowing, the negative diversity effect of undersown richness on barley disease remained significant (p < 0.001, Table S5-S6, model 4). The two species that increased barley disease when present, were *Phleum pratense* (probability of >10% leaf area damage increased from 55.31% to 63.74%, p<0.001) and *Lolium multiflorum* (probability of >10% leaf area damage increased from 54.28% to 67.79%, p=0.027, Table S5-S6, Figure S4, model 4, H1c). When we corrected for single species abundances, undersown richness had no significant effect on barley disease (p = 0.180). Instead, with increasing abundance of five out of the eight species, all of the nitrogen fixing species and *Cichorium intybus*, barley disease significantly decreased, while the abundance of the other three species had no significant effect (Table S7-S8, Figure 3, model 5, H1c).

**Figure 3:**
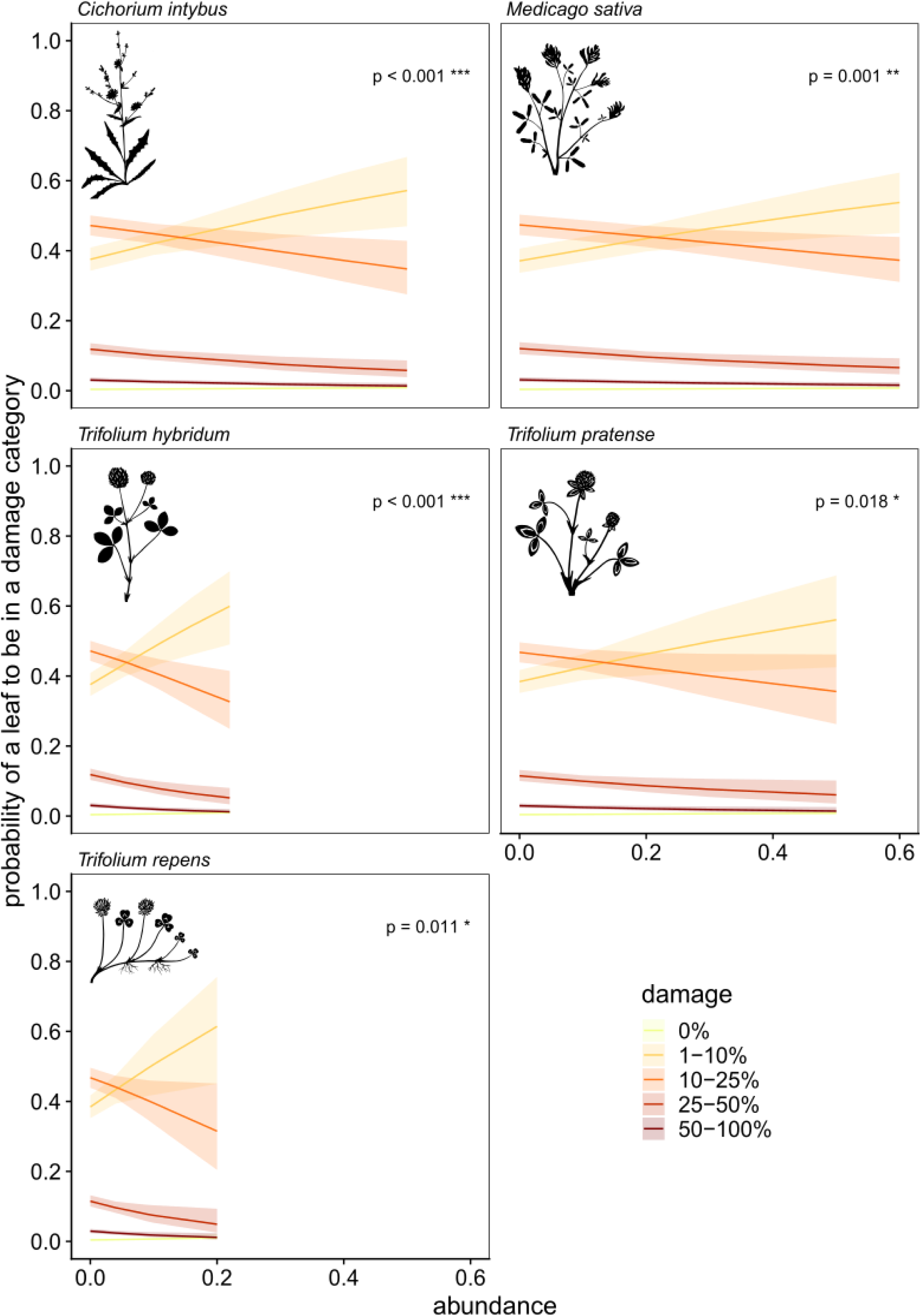
The probability of a barley leaf to fall into different damage categories as a function of the abundance of *Trifolium repens, Trifolium hybridum*, *Medicago sativa*, *Trifolium pratense* and *Cichorium intybus*, predicted by the simplified cumulative link mixed model (model 5). The fitted lines match the realized abundances of the species and are given with 95% confidence limits.

**Barley yield** was on average 2.246±0.580tha^-1^. Undersown richness explained yield better than undersown functional diversity (Table S9) and the following models therefore only include undersown richness. Undersown richness reduced yield from 2.422tha^-1^ in the absence of undersowns to 1.952tha^-1^ when all eight species were undersown, but this was only marginally significant (p=0.073, Table S10-S11, Figure S5, model 6, H3). The diversity effect was largely explained by herbicide and undersowing (Table S10-S11, model 7-8). The model analyzing yield in response to a categorical treatment variable explained more variance in yield (R^2^=0.527, model 9) than the models analyzing it in response to undersown richness (R^2^=0.227, model 6). There was one undersown species that had a large and statistically significant negative impact on yield: undersowing with *Medicago sativa* reduced yield by 50% compared with barley growing alone with (-54%) or without (-49%) herbicide (Table S12, Figure S6, model 9). The other undersowing treatments had smaller impacts on yield: undersowing with *Trifolium hybridum* (+6% compared with barley alone, -3% compared with barley with herbicide), *Trifolium repens* (+4%, -5%), *Trifolium pratense* (-11%, -19%), *Lolium multiflorum* (-11%, -19%), *Phleum pratense* (-13%, -21%), *Festuca arundinacea* (+7%, -3%), *Cichorium intybus* (-24%, -31%), two species (-11%, -17%), four species (-23%, -30%) and eight species (-9%, -17%) but none of those differences were statistically significant (Table S12, Figure S6, model 9, H3b).

### Soil transplant experiment

In the soil transplant experiment, leaves were less likely to be damaged by **disease** in early July when the pots were placed into a more species-rich environment (p = 0.012, Figure 4a). Soil-origin richness had the opposite effect, but this was not statistically significant (p = 0.43, Figure 4b). The interaction between environment and soil-origin richness was not statistically significant (p = 0.44, Table S13-S14, model 10a). In early August, none of the treatments had a statistically significant effect on disease (Table S13-S14, model 10b, H1a-b).

**Figure 4:**
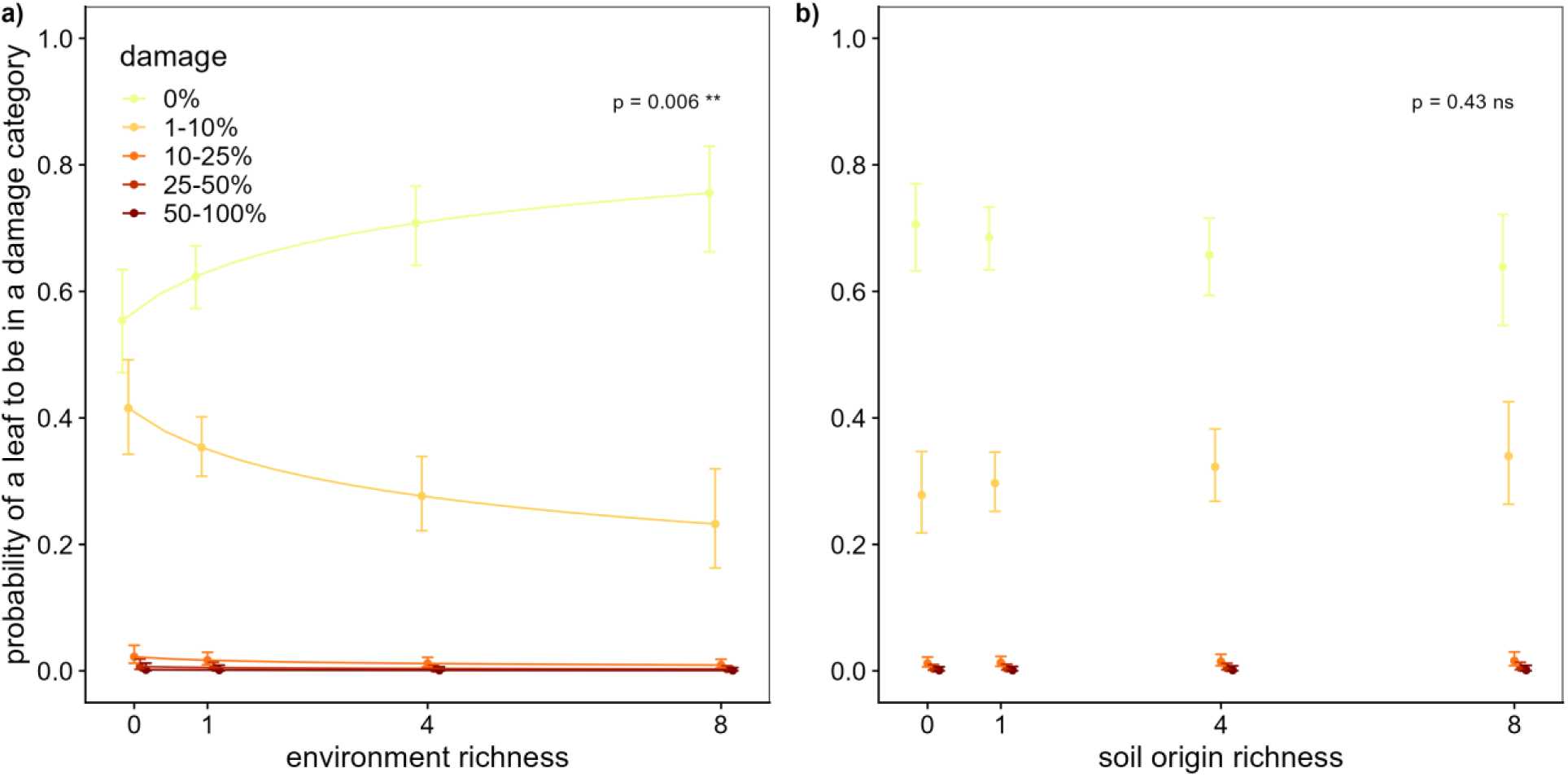
The probability of a barley leaf in the soil transplant experiment to fall into a damage category with a) increasing environment richness and b) soil-origin richness in July 2020

The **Shannon diversity of the fungal OTUs** in the pot soils increased marginally with soil-origin richness, but only when the pots were placed in a plot with high environment richness (p = 0.084, Table S15-S16, Figure S7, H2). The **proportion of potentially pathogenic OTUs** decreased both with soil origin (p < 0.001, Table S17-S18, Figure S8a) and environment richness (p = 0.039, Table S17-S18, Figure S8b), but the **relative abundance of potentially pathogenic OTU reads** was not affected by soil origin (p = 0.40, Table S19-S20, Figure S9a) or environment richness (p = 0.98, Table S19-S20, Figure S9b, H2). The **probability for the occurrence of potentially symbiotic OTUs** in the pot greatly increased with soil-origin richness (p < 0.001, Table S21-S22, Figure S10a) but did not change with environment richness (p = 0.69, Table S21-S22, Figure S10b, H2).

## Discussion

Here, we tested whether undersown plant species can be used as a sustainable disease control in an agroecosystem. We observed reduced barley foliar disease in undersown plots of the TWINWIN experiment as predicted by H1 (Figure S2 and S3). The reduced foliar disease in the pot experiment could only be attributed to environment, but not soil origin diversity (Figure 4), suggesting that, and as predicted by H1a, aboveground mechanisms exclusively drive disease in this system. Single undersown species varied in their contribution to disease reduction in the TWINWIN experiment (Figure 3), consistent with H1c, and the presence of two species was even attributed to increased disease. Later in the season, all barley leaves in both experiments were heavily diseased, independent of the treatments. Our results highlight that the addition of few undersown species can promote desired ecosystem functions and enhance sustainable agriculture, but the outcome depends on the identity of the added species.

Undersowing decreased barley foliar disease early in the season and this effect became - although only marginally significant-stronger with increasing undersown richness (Figure S2 and S3), as predicted by H1. Biodiversity experiments (X. Liu et al., 2016; Mitchell et al., 2003; Rottstock et al., 2014) and intercropping studies Kunelius et al. (1992) generally find similar effects of diversity. Undersown richness was the better predictor of barley disease than undersown functional diversity, which is also consistent with findings from a grassland experiment (Rottstock et al., 2014). However, the root traits used to create the gradient of functional diversity were not primarily chosen to optimize foliar disease reduction. Diversifying traits linked to defense against pathogens could still be a promising approach (Garrett et al., 2009). The effect of undersown diversity was explained mostly by undersowing per se, meaning that undersowing single or only few species can provide adequate disease reduction. This is consistent with ecological theory and the results from diversity experiments indicating largest benefits of diversification at low diversity (Cardinale et al., 2012; Isbell et al., 2017). The need for only few species should simplify the adoption of undersowing in agroecosystems.

Most of the species that reduced barley disease were nitrogen-fixing legumes (Figure 3), which is consistent with observations from intercropping studies (Hauggaard-Nielsen et al., 2008; Zhang et al., 2019). Legumes in crop rotations and crop mixtures enhance the nutritional status of the other crops Boetzl et al. (2023). Nitrogen fertilization has been linked to reduced pressure by certain diseases (Dordas, 2008; Tripathi et al., 2022), which may explain the reducing effects of the undersown legumes on barley foliar disease. However, the missing soil effects in our soil transplant experiment do not support such soil-mediated mechanisms. Aboveground mechanisms, like physical interference with growth and dispersal are therefore more likely to explain the disease reduction by the non-grass undersowns. Canopy structure, especially density, has been shown to reduce spore dispersal by rain splashes (Schoeny et al., 2008; Vidal et al., 2017) which is how *Phyrenophora teres* ascospores disperse (Z. Liu et al., 2011). The results of Rottstock et al. (2014) suggest canopy density and improved space filling are key components limiting spore dispersal with increasing plant diversity. Our results suggest, that leaf shape might substantially contribute to this, too. It is possible that the thin narrow leaves with upward orientation of grasses are not as well suited to capture dispersing spores as the broader leave shapes of the forbs.

Furthermore, the forbs which are only distantly related to barley, might have exposed the barley plants to microbes that are not able to successfully infect barley, as most fungal pathogens infect only closely related hosts (Klenke, 2015). However, those microbes, non-pathogenic for barley, might still have challenged the barleys immune system, leading to increased defense against *Pyrenophora teres*, as has been observed by Lyngs Jørgensen et al. (1998). In contrast, closely related neighbors can contribute to disease risk, by increasing the number of potential hosts (Parker et al., 2015). *Pyrenophora teres* can also infect *Phleum pratense* and *Lolium multiflorum* (Farr & Rossman, 22.05.2023). The stubble born pathogen *P. teres* (Z. Liu et al., 2011) might overwinter on the alternative host, which could explain why *P. pratense* and *L. multiflorum* presence increased barley foliar disease (Figure 4).

Impacts of single undersown species on yield corresponding to our study have been detected in a meta-analyses of other Nordic studies, where undersown legumes increased yield by 6% but non-legumes decreased it by 3% (Valkama et al., 2015). However, our results regarding the yield-reducing impacts of *Medicago sativa* (a legume not included Valkama et al., 2015) contrast with results from other legumes, indicating that not all legumes promote yield, as was suggested also earlier by (Abdalla et al., 2019). *M. sativa* visibly outgrew barley in many plots where it was present. Yield losses with increasing undersown richness and in plots where barley was grown with *M. sativa* suggest that mechanisms such as competition between barley and undersown species may overrule potential yield gains through disease suppression. To fully disentangle the competition-related undersowing effects from those related to disease suppression, a fungicide treatment could be included in future studies.

This study showed that undersown species can suppress crop diseases similarly as intercropping (Finckh et al., 2000; Hauggaard-Nielsen et al., 2008; Trenbath, 1993) but only early in the season and without large impact on yield. The variety of barley used here is susceptible to net blotch (Jalli & Vuorinen, 2007; Ohralahti, April 2013) which might explain why we did not detect reduction effects of undersowing later in the season. Undersowing might be able to contribute to prolonged disease suppression with positive impacts on yield if it were combined with additional disease control measures, like using resistant cultivars. The shallow rooting legumes *Trifolium repens* and *T. hybridum* reduced disease most strongly and yield was, although not statistically significant, higher in their presence compared to the barley monocultures. Those two species might also have other beneficial effects: Boetzl et al. (2023) showed that undersown clover mixtures increased pollinator abundance and suppressed weeds.

Contrary to our expectations (H1b), only environmental richness and not soil-origin richness had a statistically significant effect on foliar disease in the pot experiment (Figure 4), suggesting that aboveground mechanisms contributed to disease reduction exclusively in this study system. Processes like physical spore interception aboveground (Vidal et al., 2017), microclimatic conditions favoring barley health (Zhu et al., 2000) or allelopathic effects (Gómez-Rodríguez et al., 2003; Wenda-Piesik et al., 2010) must have been more important drivers of disease than belowground mechanisms, like improved nutritional status (Bedoussac et al., 2015), root allelopathy (Fiers et al., 2013; Kaddes et al., 2016) or effects through changes in the soil microbial community (Moya et al., 2018; Pozo & Azcón-Aguilar, 2007). However, we did not evaluate belowground disease symptoms. Future studies could focus more on belowground diseases and mechanisms driving them.

We found a shift towards a soil fungal community that should promote barley health with increasing soil-origin richness: OTU richness (Figure S7) and the presence of potentially symbiotic fungi (Figure S10) increased while the proportion of potentially pathogenic fungi decreased (Figure S8). Only the the relative abundance of potentially pathogenic OTUs did not changed with soil-origin richness. Despite this and the generally acknowledged positive effects of soil microbial diversity for plant health (Bollmann-Giolai et al., 2022; Garbeva et al., 2004), soil-origin richness did not significantly reduce foliar disease in the pot experiment. Focusing on symptoms specific to soilborne diseases might have revealed different patterns (Zhang et al., 2019), especially since we observed a lower proportion of potentially pathogenic OTUs at higher undersown richness. Furthermore, diversity effects in experiments commonly become stronger over time (e.g. Huang et al., 2018; Lange et al., 2023; Meyer et al., 2016; Reich et al., 2012) as soil microbial communities build up over years (Morriën et al., 2017). The observed undersown diversity related soil microbial community differences might affect aboveground disease symptoms only after a longer experiment duration.

## Conclusions

Undersowing barley with few effective species can reduce disease pressure via aboveground mechanisms, and hence can provide an alternative pest control to pesticides. Our study did not provide evidence of disease suppression via belowground mechanisms, but it should be noted that these may only manifest after a time lag. Jointly, our results show that agroecosystem functioning can be improved without trade-offs on yield by targeted selection of undersown species. Establishing direct links between diversification schemes, microbial responses (both above and belowground) and yield across different environmental conditions and crops offers an exciting future venue of research, and is needed to guide sustainable agricultural practices.

## Supporting information

Supplementary Methods and Results

## Acknowledgements

We thank Alma Oksanen and Heidi Blom for all their help with the experiments. We are grateful to the research farm staff for maintaining the TWINWIN experiment. Funding was provided by the Nessling Foundation and the Academy of Finland (STN MULTA; 327222) to A-L.L, A.L. and J.H.

