## Supplementary Methods and Results for "The potential of undersown species identity vs. diversity to manage disease in crops"

#### TWINWIN experiment

| Control | | Barley + 1 | | Barley + 2 | | Barley + 4 | | Barley + 8 | |
|---|---|---|---|---|---|---|---|---|---|
| Bare | x 4 | IR | x 3 | IR TG | x 1 | IR AC AA FA | x 1 | IR TG AC WC AA RC FA CI | x 4 |
| | | TG | x 3 | AC WC | x 1 | TG WC RC CI | x 1 | | |
| Barley monocult. | x 8 | AC | x 3 | AA RC | x 1 | IR TG AA FA | x 1 | | |
| | | WC | x 3 | FA CI | x 1 | AC WC RC CI | x 1 | | |
| Barley + herbicide | x 4 | AA | x 3 | AA AC | x 1 | AA RC FA CI | x 1 | | |
| | | RC | x 3 | RC FA | x 1 | IR TG AC WC | x 1 | | |
| | | FA | x 3 | AA IR | x 1 | | | | |
| | | CI | x 3 | AC FA | x 1 | | | | |
| | | | | WC IR | x 1 | | | | |
| | | | | CI TG | x 1 | | | | |

Table S1: TWINWIN design. The left side of the columns indicate a treatment, the right side of the column indicates how many plots with this treatment existed. The treatments are barley alone, barley with herbicide and barley with undersown species (B + 1, 2, 4, 8). For the latter, the treatment corresponds with the identity of the undersown species, which are *Lolium multiflorum* (Italian ray grass, IR), *Phleum pratense* (Timothy grass, TG), *Trifolium hybridum* (Alsike clover, AC), *Trifolium repens* (White clover, WC), *Medicago sativa* (Alfalfa, AA), *Trifolium pratense* (Red clover, RC), *Festuca arundinacea* (Tall fescue, FA) and *Cichorium intybus* (common chicory, CI). The functional traits of the species are described in the methods. Not all theoretically possible species combinations were realized in the barley + 2 and barley + 4 treatments.

Cover measurements:

We visually assessed the percent cover of all sown plant species separately, all weeds together and bare ground in five 1m2 subplots per plot end of July 2020. Weed cover was measured as quality control and the relative abundance of weeds was below 5% in all plots. The sum of all cover values per subplot could exceed 100%, due to layered vegetation structure. We transformed the values per subplot to **relative abundances** and averaged them per plot and species. We standardized the assessments done by multiple people, and everybody evenly analyzed subplots in every plot to minimize bias.

#### Soil transplant experiment

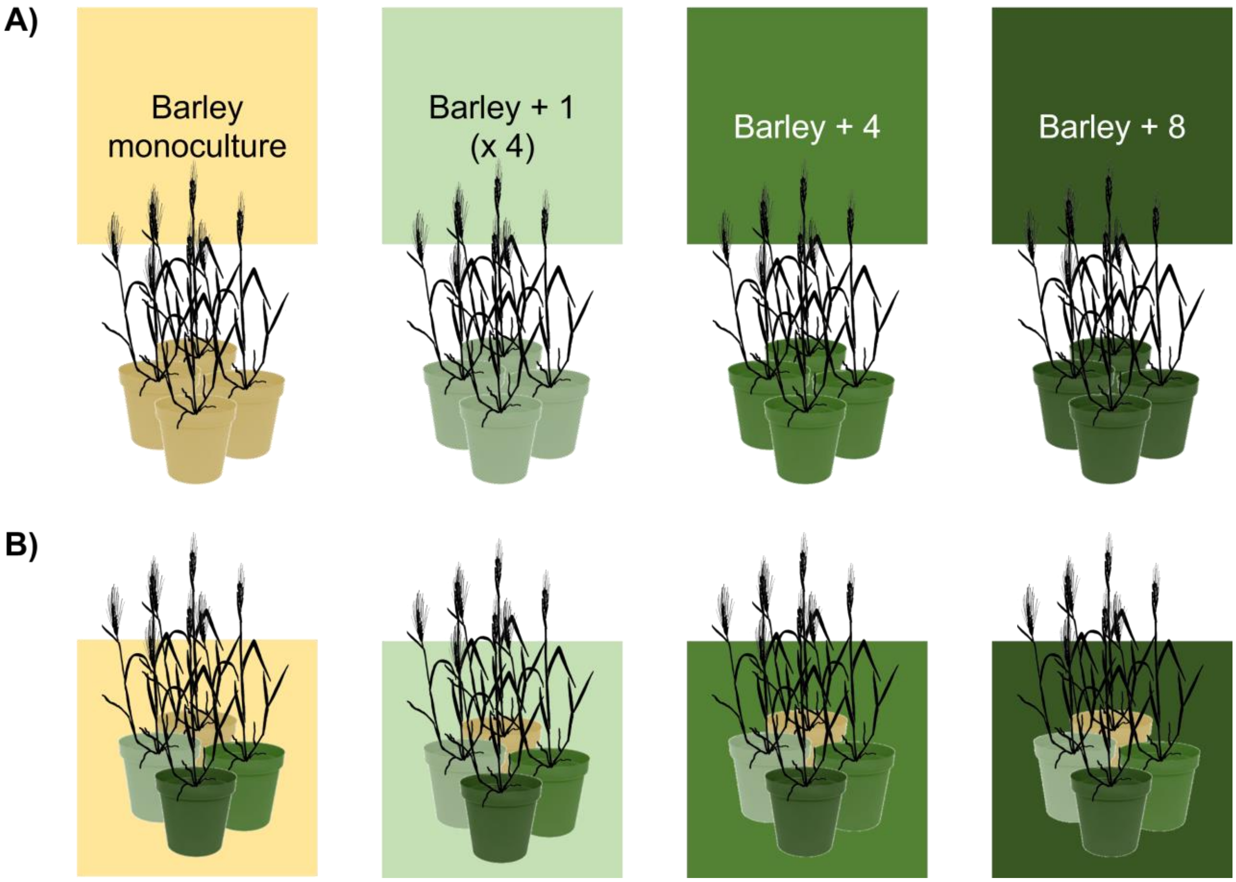


Figure S1: Soil transplant experiment. A) Barley was grown in pots with soil from plots varying in undersown richness (soil-origin richness) and B) placed reciprocally back to the plots where the soil originated from (environment richness). For this experiment a subset of all plots was used: all eight barley monocultures, all twelve barley plots which grew with one of the following species: *Trifolium hybridum*, *Medicago sativa*, *Lolium multiflorum* or *Festuca arundinacea*, the single plot in which all of these four species were grown together with barley, and the four plots in which barley was grown with all eight undersown species.

Set up of pots:

Three barley seeds were germinated and grown in well-watered, sand-filled cardboard pots, in the greenhouse in the beginning of June 2020. After two weeks, the seedlings were transplanted with the cardboard pots into 12 x 12 cm pots of 20 cm height filled with a 0.5 L layer of sand at the bottom and a 1l 1:3 mixture of field soil (soil origin according to the design described in Supplementary methods) and sand layer. The field soil was freshly collected from the TWINWIN plots and soil from plots with identical undersowing treatment was mixed. We added a 1 cm layer of sand on top to suppress weed germination from the seedbank in the field soil and to reduce water evaporation. Not all seeds germinated, which is why we removed seedlings so that each pot contained two individuals. In the last week of June 2020, all pots were moved to the field and placed in the plots between the barley rows. When there were multiple plots with the same undersown community, we randomly assigned the pots to one of the plots. Each pot was placed in an individual plastic tray to minimize contact with the soil organisms of their surroundings. Pots received water ad libitum at the beginning of the experiment and during the driest periods of the growing season. An additional tablespoon of newly collected soil was added to the pots in mid July 2020 to renew the inoculum. The additional soil was collected in the same plots as the originally used soil. Plants in one pot in the undersown richness 4 plot were destroyed by birds during the experiment.

Soil microbial analyses

We collected soil samples from the pots when the barley was ready for harvest early September 2020. These soil samples were frozen at -20°C to wait for DNA extraction. For practical reasons, we used only subset of all pots for this, which was chosen so that we had equal number of pots per soil origin treatment (33, total 231 samples), and at least three pots from each environment x soil combination.

DNA extraction was done using Zymo Quick-DNA Soil Microbe 96 Kit according to manufacturer instructions. Technical duplicates of DNA extraction were pooled before quantification using Qubit to improve reproducibility. We used the ITS sequencing region (ITS7 - ITS4, Ihrmark et al. (2012)) tagged amplicon sequencing on the NovaSeq PE250 platform by Novogene (Cambrigde, UK). Raw sequences from amplicon sequencing were quality filtered, merged, and clustered to generate OTU's at 97% sequence similarity using QIIME2 pipeline (Bolyen et al., 2019). UNITE (version 01.12.2017, (Kõljalg et al., 2005) was used for the fungal taxonomy assignment. We excluded all OTUs that had no blast hit (4143 out of total 8095 OTUs) or that were not classified as fungi (270 OTUs) to be sure we only included fungal taxa.

To characterize the soil fungal communities in the pots that differed in soil-origin richness and environment richness we calculated a set of metrics, related to OTU diversity and the presence and abundance of potentially symbiotic and pathogenic fungal taxa for each pot: To assess fungal OTU diversity, we calculated the Shannon diversity index using number of reads per OTU as measure of abundance. We used the FUNGuild database to assess for every OTU whether was is potentially plant pathogenic or whether it was an arbuscular mycorrhizal fungus and therefore potentially symbiotic (Nguyen et al., 2016). FUNGuild assignes fungal taxa into 12 categories: animal pathogens, arbuscular mycorrhizal fungi, ectomycorrhizal fungi, ericoid mycorrhizal fungi, foliar endophytes, lichenicolous fungi, lichenized fungi, mycoparasites, plant pathogens, undefined root endophytes, undefined saprotrophs, and wood saprotrophs. Every OTU that could potentially be a plant pathogen was classified as potentially pathogenic. Any OTU that was classified as arbuscular mycorrhizal fungi was classified as potentially symbiotic. We then calculated the relative abundance of potentially pathogenic OTU reads as the proportion of reads of all potentially pathogenic OTUs out of all OTU reads with a guild assigned in a pot. To better account for rare OTUs we calculated the proportion of potentially pathogenic OTUs from all the OTUs with a guild assigned in funguild. We calculated the relative abundance of potentially symbiotic OTU reads and the proportion of potentially symbiotic OTUs the same way as for potentially pathogenic OTUs. Only 38% of all pots contained OTUs that were classified as potentially symbiotic OTUs and when they were present, their relative abundance was very low (<0.15%). Because of this, we only looked at the presence/absence of potentially symbiotic OTUs.

### Supplementary Results

#### TWINWIN disease models

| model | df | AIC |
|---|---|---|
| disease ~ block + leaf + log(richness + 1) * functional diversity | 13 | 6,983.070 |
| disease ~ block + leaf + log(richness + 1) + functional diversity | 12 | 6,984.901 |
| disease ~ block + leaf + log(richness + 1) | 11 | 6,983.319 |
| disease ~ block + leaf + functional diversity | 11 | 6,984.332 |

Table S2: AIC comparisons for four cumulative link mixed effects models including variable combinations of functional diversity and undersown richness to explain disease in the TWINWN experiment in July 2020.

| | 1. richness model | | | 2. herbicide model | | | 3. undersowing model | | |
|---|---|---|---|---|---|---|---|---|---|
| Variable | Chi2 | P-value | | Chi2 | P-value | | Chi2 | P-value | |
| block | 25.36 | < 0.001 | *** | 25.74 | < 0.001 | *** | 28.44 | < 0.001 | *** |
| leaf | 2,197.21 | < 0.001 | *** | 2,197.36 | < 0.001 | *** | 2,199.52 | < 0.001 | *** |
| herbicide | | | | 1.07 | 0.301 | | 4.32 | 0.038 | * |
| undersowing | | | | | | | 6.95 | 0.008 | ** |
| log richness | 3.35 | 0.067 | . | 4.36 | 0.037 | * | 0.00 | 0.961 | |

Table S3: Anova tables to test for the overall effect of explanatory variables for the three cumulative link mixed effects models that analyze disease in the TWINWIN experiment in July 2020 with block, leaf position and log-transformed undersown richness as fixed effects and plot identity and unique plant individual as random effect. Model 2 and 3 test if herbicide and undersowing explain the diversity effect observed in model 1.

| | 1. richness model | | | | 2. herbicide model | | | | 3. undersowing model | | | |
|---|---|---|---|---|---|---|---|---|---|---|---|---|
| Fixed effects | Estimate | SD | z-value | P-value | Estimate | SE | z-value | P-value | Estimate | SE | z-value | P-value |
| block2 | -0.0877 | 0.173 | -0.51 | 0.612 | -0.0864 | 0.1714 | -0.5 | 0.614 | -0.0956 | 0.161 | -0.59 | 0.553 |
| block3 | 0.5748 | 0.1728 | 3.33 | 0.001 *** | 0.5742 | 0.1711 | 3.36 | 0.001 *** | 0.5739 | 0.1608 | 3.57 | < 0.001 *** |
| block4 | 0.7001 | 0.173 | 4.05 | < 0.001 *** | 0.701 | 0.1713 | 4.09 | < 0.001 *** | 0.6926 | 0.1609 | 4.3 | < 0.001 *** |
| leaf position | -2.6971 | 0.0754 | -35.76 | < 0.001 *** | -2.6971 | 0.0754 | -35.76 | < 0.001 *** | -2.6982 | 0.0754 | -35.78 | < 0.001 *** |
| herbicide | | | | | -0.2613 | 0.2514 | -1.04 | 0.299 | -0.547 | 0.2583 | -2.12 | 0.034 * |
| undersowing | | | | | | | | | -0.582 | 0.2141 | -2.72 | 0.007 ** |
| log richness | -0.1859 | 0.1003 | -1.85 | 0.064 . | -0.228 | 0.1073 | -2.13 | 0.034 * | 0.0053 | 0.132 | 0.04 | 0.968 |
| Random effects | Variance | SD | | | Variance | SD | | | Variance | SD | | |
| individual ID | 1.4026 | 1.1843 | | | 1.4021 | 1.1841 | | | 1.4078 | 1.1865 | | |
| plot | 0.0543 | 0.2331 | | | 0.0504 | 0.2244 | | | 0.0259 | 0.1609 | | |
| Threshold coefficients | Estimate | SE | z-value | | Estimate | SE | z-value | | Estimate | SE | z-value | |
| 0\|1 | -10.6552 | 0.3 | -35.52 | | -10.7079 | 0.3045 | -35.17 | | -11.0019 | 0.3236 | -34 | |
| 1\|2 | -5.6161 | 0.2152 | -26.09 | | -5.6692 | 0.2209 | -25.66 | | -5.9624 | 0.2434 | -24.49 | |
| 2\|3 | -3.3806 | 0.1881 | -17.97 | | -3.434 | 0.1944 | -17.66 | | -3.726 | 0.218 | -17.09 | |
| 3\|4 | -1.6655 | 0.1777 | -9.38 | | -1.7188 | 0.184 | -9.34 | | -2.0091 | 0.2074 | -9.69 | |

Table S4: Summary table for the basic cumulative link mixed-effects models analyzing TWINWIN disease in July 2020 (model 1, Cond. H = 340.) and the models including herbicide (model 2. Cond. H = 360) and herbicide and undersowing (model 3, Cond. H = 540)

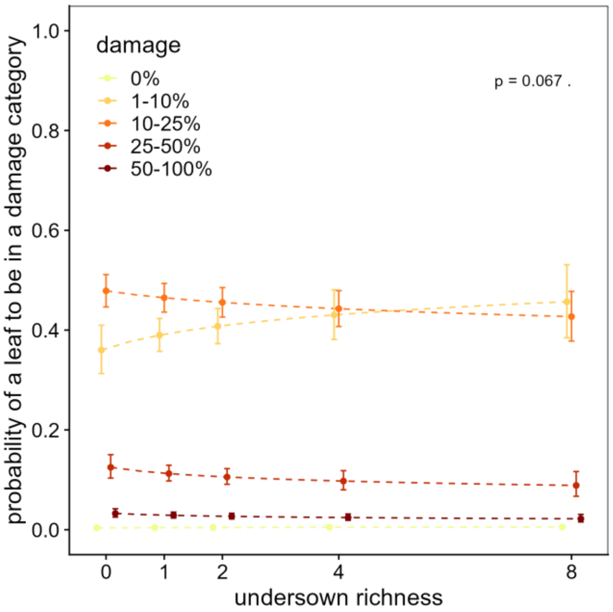


Figure S2: The predicted probability and the 95% confidence interval of a leaf to fall into a given damage category in July 2020 depending on the undersown richness based on the basic cumulative link mixed effects model, not correcting for herbicide and undersowing (model 1). Significant effects are shown as solid lines, marginally significant effects in dashed lines and for non-significant effects the model predictions are shown without a line connecting them.

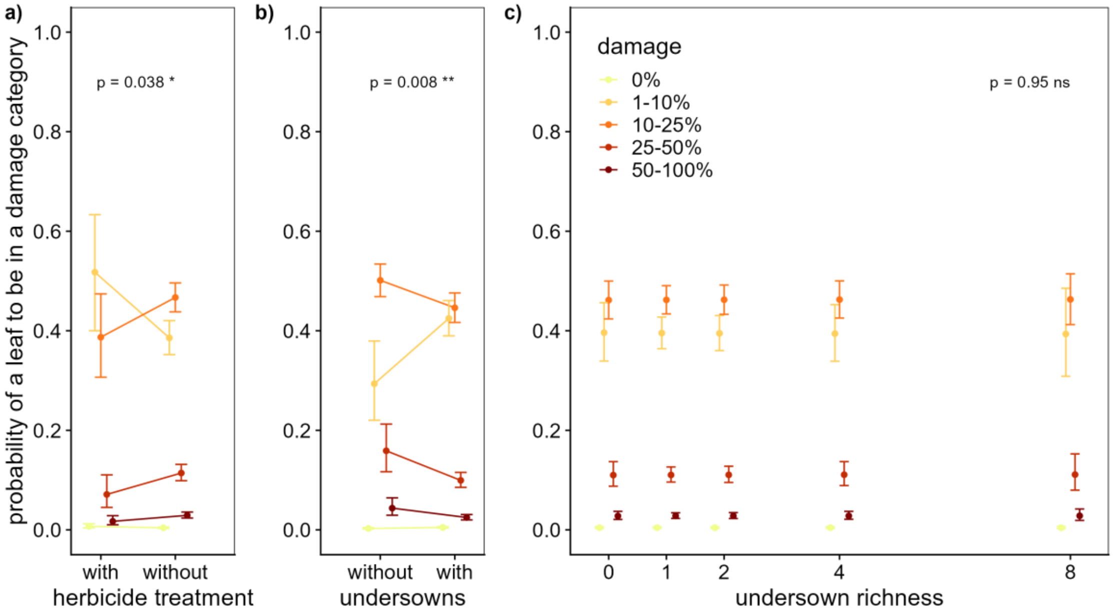


Figure S3: The predicted probability and the 95% confidence interval of a leaf to fall into a given damage category in July 2020 depending on a) herbicide treatment, b) undersowing and c) undersown richness based on the cumulative link mixed effects model including herbicide and undersowing. Significant effects are shown as solid lines, marginally significant effects in dashed lines and for non-significant effects the model predictions are shown without a line connecting them.

| Variable | Chi2 | P-value | |
|---|---|---|---|
| block | 30.90 | < 0.001 | *** |
| leaf position | 2,200.00 | < 0.001 | *** |
| log richness | 18.50 | < 0.001 | *** |
| Phleum pratense presence | 11.80 | < 0.001 | *** |
| Lolium multiflorum presence | 4.91 | 0.027 | * |
| Festuca arundinacea presence | 0.94 | 0.33 | |
| Trifolium pratense presence | 0.82 | 0.37 | |
| Trifolium repens presence | 0.71 | 0.4 | |
| Medicago sativa presence | 0.56 | 0.56 | |
| Cichorium intybus presence | 0.20 | 0.65 | |
| Trifolium hybridum presence | 0.00 | 0.99 | |

Table S5: Model selection table for the cumulative link mixed effects model to explain the proportion of leaves in different disease categories in the TWINWIN experiment plots. The full model included the presence/absence of the undersown species, undersown species richness, block and leaf position as fixed effect. Plot identity and the ID of the plant individual from which a leaf originated was included as a random effects (model 4). Anova was used to compare the models. Species presence/absence terms that did not improve the model fit were stepwise removed.

| Fixed effects | Estimate | SE | z-value | P-value | |
|---|---|---|---|---|---|
| block2 | -0.0253 | 0.1509 | -0.17 | 0.867 | |
| block3 | 0.6402 | 0.1495 | 4.28 | < 0.001 | *** |
| block4 | 0.6709 | 0.1491 | 4.5 | < 0.001 | *** |
| leaf position | -2.6969 | 0.0754 | -35.77 | < 0.001 | *** |
| log richness | -0.552 | 0.1196 | -4.62 | < 0.001 | *** |
| P. pratense presence | 0.3507 | 0.156 | 2.25 | 0.025 | * |
| L. multiflorum presence | 0.5728 | 0.1597 | 3.59 | < 0.001 | *** |
| Random effects | Variance | SD | | | |
| individual ID | 1.4 | 1.18 | | | |
| plot | < 0.0001 | < 0.0001 | | | |
| Threshold coefficients | Estimate | SE | z-value | | |
| 0\|1 | -10.727 | 0.292 | -36.7 | | |
| 1\|2 | -5.688 | 0.203 | -28 | | |
| 2\|3 | -3.452 | 0.174 | -19.8 | | |
| 3\|4 | -1.735 | 0.162 | -10.7 | | |

Table S6: Summary table for the simplified cumulative link mixed-effects model analyzing TWINWIN disease in July 2020 including presence/absence data of single undersown species (simplified model 4). Cond. H = 380

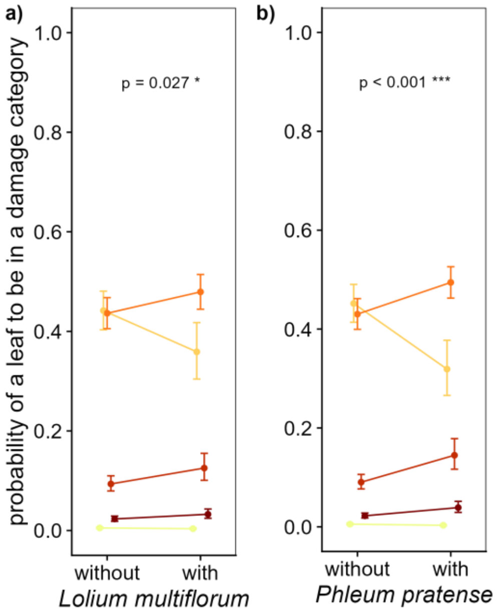


Figure S4: The predicted probability and the 95% confidence interval of a leaf to fall into a given damage category in July 2020 depending on a) Lolium multiflorum presence, b) Phleum pratense presence based on the simplified cumulative link mixed effects model including presence/absence of undersown species (simplified model 4). Significant effects are shown as solid lines, marginally significant effects in dashed lines and for non-significant effects the model predictions are shown without a line connecting them. Colors indicate the different disease categories, with darker colors representing higher disease levels (see legend in Figure 3 and Figure S2

| Variable | Chi2 | P-value | |
|---|---|---|---|
| block | 20.40 | < 0.001 | *** |
| leaf position | 2,197.00 | < 0.001 | *** |
| log richness | 1.77 | 0.18 | |
| log Trifolium hybridum abundance | 12.80 | < 0.001 | *** |
| log Cichorium intybus abundance | 11.40 | < 0.001 | *** |
| log Medicago sativa abundance | 10.50 | 0.0012 | ** |
| log Trifolium repens abundance | 6.48 | 0.011 | * |
| log Trifolium pratense abundance | 5.61 | 0.018 | * |
| log Phleum pratense abundance | 1.45 | 0.23 | |
| log barley abundance | 0.12 | 0.72 | |
| log Festuca arundinacea abundance | 0.20 | 0.66 | |
| log Lolium multiflorum abundance | 0.10 | 0.76 | |
| log weeds abundance | 0.05 | 0.82 | |

Table S7: Model selection table for the cumulative link mixed effects model to explain the proportion of leaves in different disease categories in the TWINWIN experiment plots. The full model included the abundance of the undersown species, undersown species richness, block and leaf position as fixed effects. Plot identity and the ID of the plant individual from which a leaf originated was included as a random effects (model 5). Anova was used to compare the models. Species abundance terms that did not improve the model fit were stepwise removed.

| Fixed effects | Estimate | SE | z-value | P-value | |
|---|---|---|---|---|---|
| block2 | -0.0864 | 0.1538 | -0.56 | 0.574 | |
| block3 | 0.4359 | 0.1535 | 2.84 | 0.005 | ** |
| block4 | 0.5145 | 0.1592 | 3.23 | 0.001 | ** |
| leaf position | -2.6955 | 0.0754 | -35.74 | < 0.001 | *** |
| log richness | 0.1451 | 0.109 | 1.33 | 0.183 | |
| log T. hybridum abundance | -4.7085 | 1.2948 | -3.64 | < 0.001 | *** |
| log T. repens abundance | -5.2949 | 2.0785 | -2.55 | 0.011 | * |
| log M. sativa abundance | -1.4794 | 0.4521 | -3.27 | 0.001 | ** |
| log T. pratense abundance | -1.8105 | 0.7645 | -2.37 | 0.018 | * |
| log C. intybus abundance | -2.0197 | 0.5895 | -3.43 | 0.001 | *** |
| Random effects | Variance | SD | | | |
| individual ID | 39 | 1.18 | | | |
| plot | < 0.0001 | < 0.0001 | | | |
| Threshold coefficients | Estimate | SE | z-value | | |
| 0\|1 | -10.848 | 0.298 | -36.4 | | |
| 1\|2 | -5.807 | 0.21 | -27.7 | | |
| 2\|3 | -3.571 | 0.181 | -19.8 | | |
| 3\|4 | -1.855 | 0.169 | -11 | | |

Table S8: Summary table for the simplified cumulative link mixed-effects model analyzing TWINWIN disease in July 2020 including abundance data of single undersown species (simplified model 5). H = 9500.

#### TWINWIN yield models

| Model | Df | AIC |
|---|---|---|
| yield ~ block + log(richness + 1) * funct.div | 8 | 96.67 |
| yield ~ block + log(richness + 1) + funct.div | 7 | 96.34 |
| yield ~ block + funct.div | 6 | 95.07 |
| yield ~ block + richness | 6 | 94.41 |

Table S9: AIC comparisons for four linear models including variable combinations of functional diversity and undersown richness to explain yield in the TWINWN experiment in 2020.

| | 6. richness model | | 7. herbicide model | | 8. undersowing model | |
|---|---|---|---|---|---|---|
| Variable | Chi2 | P-value | Chi2 | P-value | Chi2 | P-value |
| block | 3.88 | P-Value | 3.91 | 0.014 * | 3.89 | 0.014 * |
| herbicide | - | - | 3.24 | 0.078 . | 3.23 | 0.079 . |
| undersowing | - | - | - | - | 2.13 | 0.151 |
| log richness | 3.35 | 0.073 . | 1.57 | 0.216 | 0.16 | 0.692 |

Table S10: Anova tables to test for the overall effect of explanatory variables for the three linear models that analyze yield in the TWINWIN experiment in 2020 with block, and log-transformed undersown richness as explanatory variable. Model 7 and 8 test if herbicide and undersowing explain the undersown richness effect observed in model 6.

| | | | 1. richness model | | | | 2. herbicide model | | | | 3. undersowing model | |
|---|---|---|---|---|---|---|---|---|---|---|---|---|
| Variable | Estimate | SD | z-value | P-value | Estimate | SE | z-value | P-value | Estimate | SE | z-value | P-value |
| intercept | 2.035 | 0.170 | 11.98 | < 0.001 *** | 1.964 | 0.179 | 10.97 | < 0.001 *** | 2.0772 | 0.2233 | 9.30 | < 0.001 *** |
| block2 | 0.394 | 0.200 | 1.97 | 0.054 . | 0.392 | 0.199 | 1.97 | 0.055 . | 0.3890 | 0.1998 | 1.95 | 0.057 . |
| block3 | 0.487 | 0.200 | 2.44 | 0.018 * | 0.487 | 0.199 | 2.45 | 0.018 * | 0.4874 | 0.1997 | 2.44 | 0.018 * |
| block4 | 0.665 | 0.200 | 3.32 | 0.002 ** | 0.663 | 0.199 | 3.33 | 0.002 ** | 0.6596 | 0.1998 | 3.30 | 0.002 ** |
| herbicide | | | | | 0.353 | 0.295 | 1.20 | 0.237 | 0.2418 | 0.3236 | 0.75 | 0.459 |
| undersowing | | | | | | | | | -0.2273 | 0.2669 | -0.85 | 0.399 |
| log richness | -0.214 | 0.117 | -1.83 | 0.073 . | -0.157 | 0.126 | -1.25 | 0.216 | -0.0658 | 0.1654 | -0.40 | 0.692 |

Table S11: Summary table for the basic linear model analyzing TWINWIN yield in 2020 (model 6. R2 = 0.227) and the models including herbicide (model 7. R2 = 0.249) and herbicide and undersowing (model 8. R2 = 0.260).

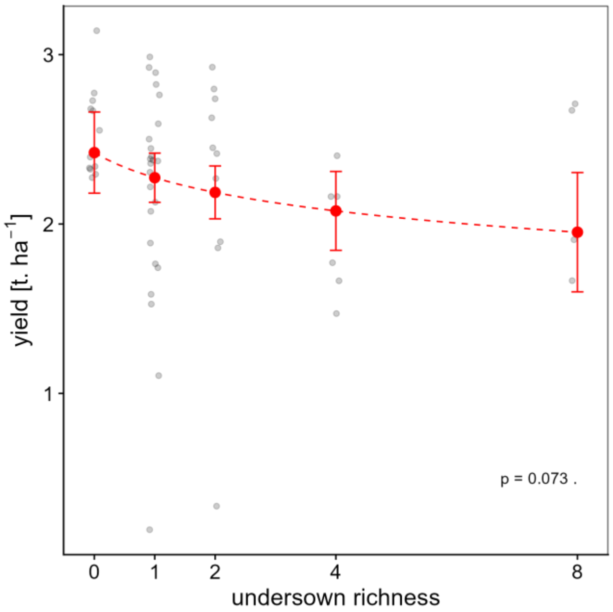


Figure S5: Yield in response to undersown richness as predicted based on the basic linear model (model 6). Raw data are in black and model prediction with the 95% interval are in red. Significant effects are shown as solid lines, marginally significant effects in dashed lines and for non-significant effects the model predictions are shown without a line connecting them.

| Variable | Chi2 | P-value |
|---|---|---|
| block | 4.97 | 0.005 ** |
| treatment | 2.48 | 0.016 * |

Table S12: Anova tables to test for the overall effect of explanatory variables for the linear model that analyzes yield in the TWINWIN experiment in 2020 with block and a categorical treatment variable that distinguishes between barley plus herbicide, barley alone, barley plus any of the undersown species alone, barley plus 2, 4 and 8 undersown species, as explanatory variable. R2 = 0.527.

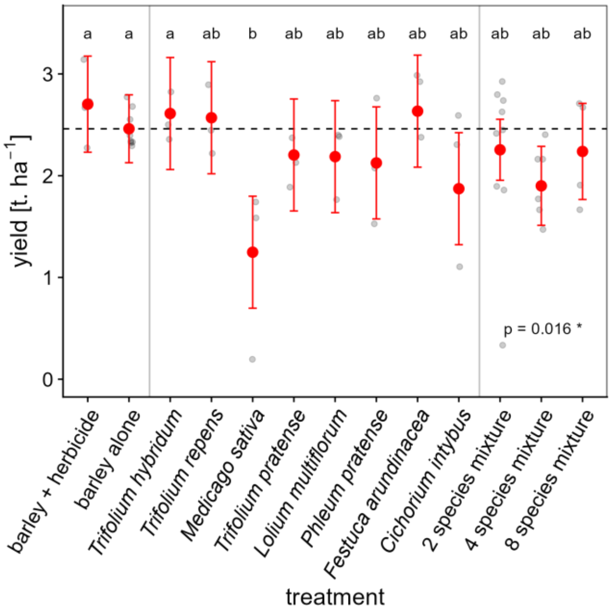


Figure S6: Yield in response to undersown treatment as predicted based on the linear model that included a categorical treatment variable instead of undersown richness (model 9). Raw data are in black and model prediction with the 95% interval are in red). The horizontal dashed line is at 2.46 t.ha-1, which equals the yield in the barley alone plots. Significance of pairwise differences was obtained with tukey-HSD post-hoc test and are indicated with letters. Estimates assigned the same letter do not statistically differ from each other.

#### Soil transplant pot experiment disease models

| | 10a. Early July 2020 | | | 10b. Early August 2020 | | |
|---|---|---|---|---|---|---|
| Variable | Chi2 | P-value | | Chi2 | P-value | |
| leaf position | 46.50 | < 0.001 | *** | 447.00 | < 0.001 | *** |
| log environment richness | 7.69 | 0.006 | ** | 0.07 | 0.79 | |
| log soil origin richness | 0.64 | 0.42 | | 0.87 | 0.35 | |
| log e. rich. x log s.o. rich. | 0.60 | 0.44 | | 1.47 | 0.23 | |

Table S13: Anova tables to test for the overall effect of explanatory variables for the cumulative link mixed-effects models that analyze disease in the soil transplant experiment in early July 2020 and early August 2020 with leaf position, log-transformed soil-origin richness, log-transformed environment richness and their interaction as fixed effects and plot identity as random effect (model 10a and 10b).

| | 10a. Early July 2020 | | | | 10b. Early August 2020 | | | |
|---|---|---|---|---|---|---|---|---|
| Fixed effects | Estimate | SD | z-value | P-value | Estimate | SE | z-value | P-value |
| leaf position | -0.6794 | 0.1029 | -6.6 | < 0.001 *** | -2.287 | 0.1251 | -18.28 | < 0.001 *** |
| log environment richness | -0.5871 | 0.234 | -2.51 | 0.012 * | 0.2131 | 0.2045 | 1.04 | 0.3 |
| log soil origin richness | -0.0309 | 0.2026 | -0.15 | 0.879 | 0.0797 | 0.1811 | 0.44 | 0.66 |
| log e. rich. x log s.o. richness | 0.144 | 0.1855 | 0.78 | 0.438 | -0.1866 | 0.1542 | -1.21 | 0.23 |
| Random effects | Variance | SD | | | Variance | SD | | |
| plot | 0.0588 | 0.242 | | | 0.0938 | 0.306 | | |
| Threshold coefficients | Estimate | SE | z-value | | Estimate | SE | z-value | |
| 0\|1 | -1.162 | 0.319 | -3.64 | | -6.893 | 0.404 | -17.08 | |
| 1\|2 | 2.074 | 0.376 | 5.52 | | -4.719 | 0.35 | -13.49 | |
| 2\|3 | 3.382 | 0.534 | 6.33 | | -3.395 | 0.323 | -10.5 | |
| 3\|4 | 4.997 | 1.042 | 4.8 | | -1.86 | 0.304 | -6.12 | |

Table S14: Summary table for the cumulative link mixed-effects models that analyze disease in the soil transplant experiment in early July 2020 and early August 2020 with leaf position, log-transformed soil-origin richness, log-transformed environment richness and their interaction as fixed effects and plot identity as random effect (model 10a. Cond. H = 980 and 10b. cond. H = 380).

#### Soil transplant pot experiment soil microbial community models

| Variable | Chi2 | P-value |
|---|---|---|
| log environment richness | 0.74 | 0.391 |
| log soil origin richness | 0.00 | 0.957 |
| log e. rich. x log s.o. rich. | 3.02 | 0.084 . |

Table S15: Anova table to test for the overall effect of explanatory variables table for the linear mixed-effects model of OTU Shannon diversity of the soil microbial community with log-transformed soil-origin richness, log-transformed environment richness and their interaction as fixed effects and plot identity as random effect (model 11).

| Fixed effects | Estimate | SE | t-value | P-value |
|---|---|---|---|---|
| intercept | 2.8218 | 0.2829 | 9.98 | < 0.001 *** |
| log environment richness | -0.172 | 0.2002 | -0.86 | 0.391 |
| log soil origin richness | 0.0108 | 0.0108 | 0.05 | 0.957 |
| log e. rich. x log s.o. richness | 0.2465 | 0.1419 | 1.74 | 0.084 . |
| Random effects | Variance | SD | | |
| plot | 0 | 0 | | |
| residual | 0.309 | 0.556 | | |

Table S16: Summary table for the linear mixed effects model of OTU Shannon diversity of the soil microbial community with log-transformed soil-origin richness, log-transformed environment richness and their interaction as fixed effects and plot identity as random effect (model 11).

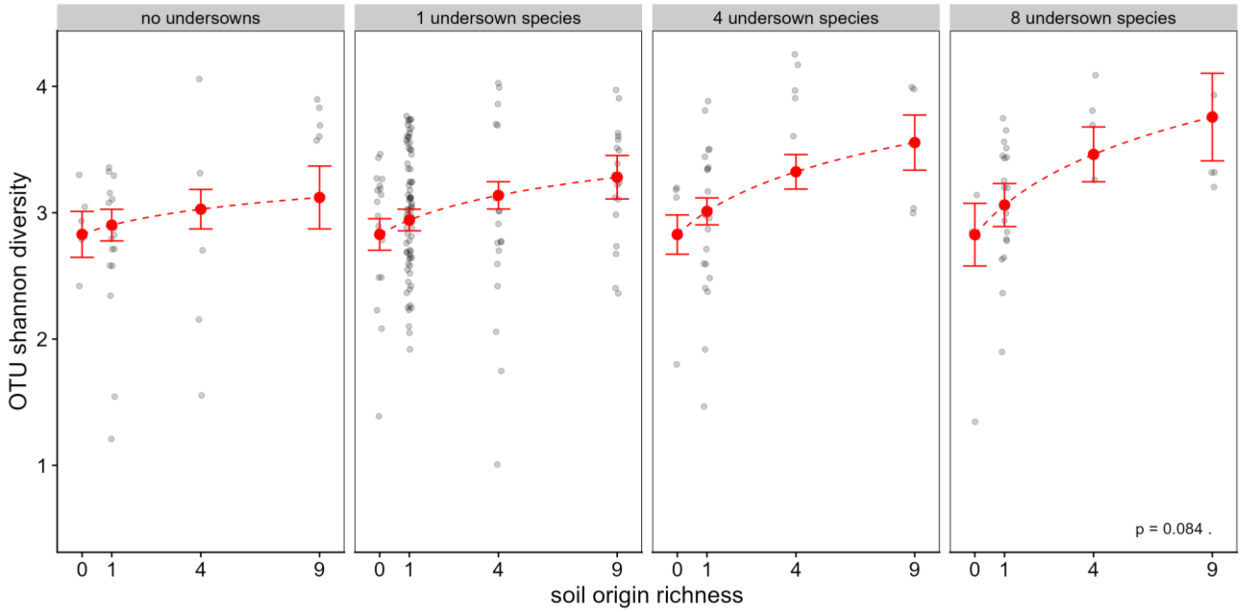


Figure S7: OTU Shannon diversity of the soil microbial community in response to soil-origin richness and environment richness. Raw data are in black and model prediction on the linear mixed effects model (model 11) with the 95% interval are in red. Marginally significant effects in dashed lines.

| Variable | Chi2 | P-value | |
|---|---|---|---|
| log environment richness | 4.25 | 0.039 | * |
| log soil origin richness | 48.10 | < 0.001 | *** |
| log e. rich. x log s.o. rich. | 0.73 | 0.39 | |

Table S17: Anova table to test for the overall effect of explanatory variables table for the generalized linear mixed-model of the proportion of potentially pathogenic OTUs with log-transformed soil-origin richness, log-transformed environment richness and their interaction as fixed effects and plot identity as random effect (model 12).

| Fixed.Effects | Estimate | SE | z-value | P-value | |
|---|---|---|---|---|---|
| intercept | -1.3519 | 0.04533 | -29.82 | < 0.001 | *** |
| log environment richness | -0.00372 | 0.03236 | -0.11 | 0.908 | |
| log soil origin richness | -0.05802 | 0.03144 | -1.85 | 0.065 | . |
| log e. rich. x log s.o. richness | -0.01915 | 0.02249 | -0.85 | 0.395 | |
| Random effects | Variance | SD | | | |
| plot | 0.000304 | 0.0174 | | | |

Table S18: Summary table for the generalized linear mixed model of the proportion of potentially pathogenic OTUs with log-transformed soil-origin richness, log-transformed environment richness and their interaction as fixed effects and plot identity as random effect (model 12).

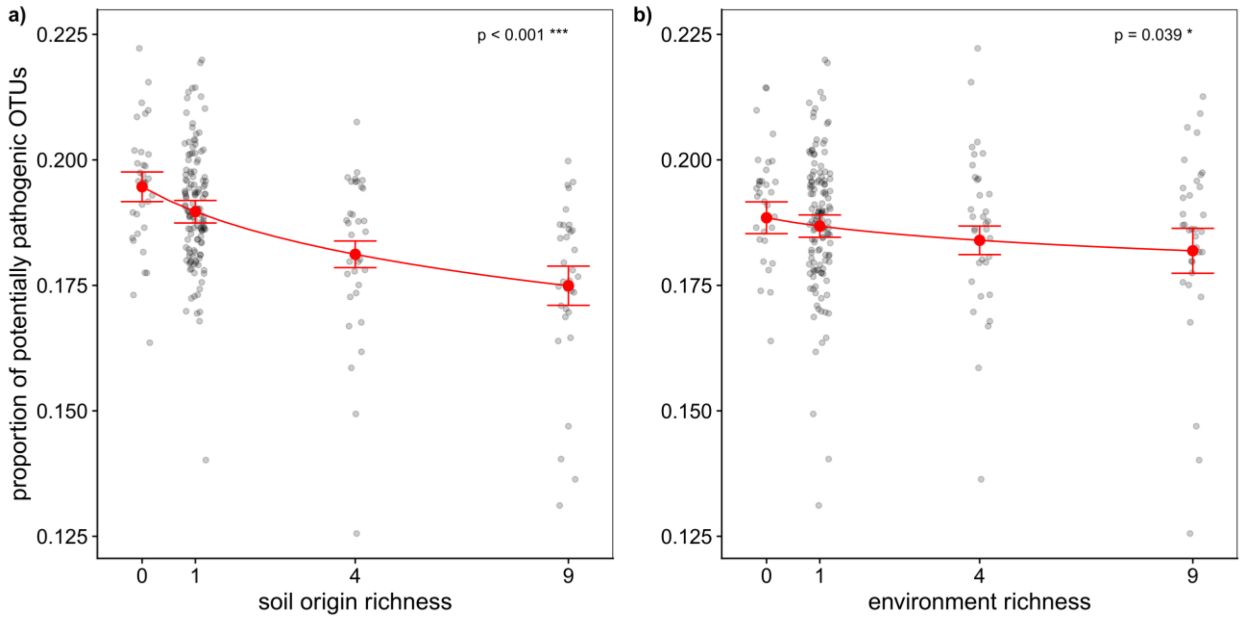


Figure S8: The proportion of potentially pathogenic fungi in the soil transplant experiment in response to a) soil-origin richness and b) environment richness. Raw data are in black and model prediction on the generalized linear mixed model (model 12) with the 95% interval are in red. Significant effects are shown as solid lines.

| Variable | Chi2 | P-value |
|---|---|---|
| log environment richness | 0.00 | 0.98 |
| log soil origin richness | 0.72 | 0.40 |
| log e. rich. x log s.o. rich. | 2.24 | 0.13 |

Table S19: Anova table to test for the overall effect of explanatory variables table for the generalized linear mixed model of the relative abundance of potentially pathogenic OTU reads with log-transformed soil-origin richness, log-transformed environment richness and their interaction as fixed effects and plot identity as random effect (model 13).

| Fixed.Effects | Estimate | SE | z-value | P-value |
|---|---|---|---|---|
| intercept | -2.147 | 0.271 | -7.91 | < 0.001 *** |
| log environment richness | -0.263 | 0.19 | -1.38 | 0.17 |
| log soil origin richness | -0.209 | 0.192 | -1.09 | 0.28 |
| log e. rich. x log s.o. richness | 0.203 | 0.133 | 1.52 | 0.13 |
| Random effects | Variance | SD | | |
| plot | 7.85E-10 | 2.80E-05 | | |

Table S20: Summary table for he generalized linear mixed model of the relative abundance of potentially pathogenic OTU reads with log-transformed soil-origin richness, log-transformed environment richness and their interaction as fixed effects and plot identity as random effect (model 13).

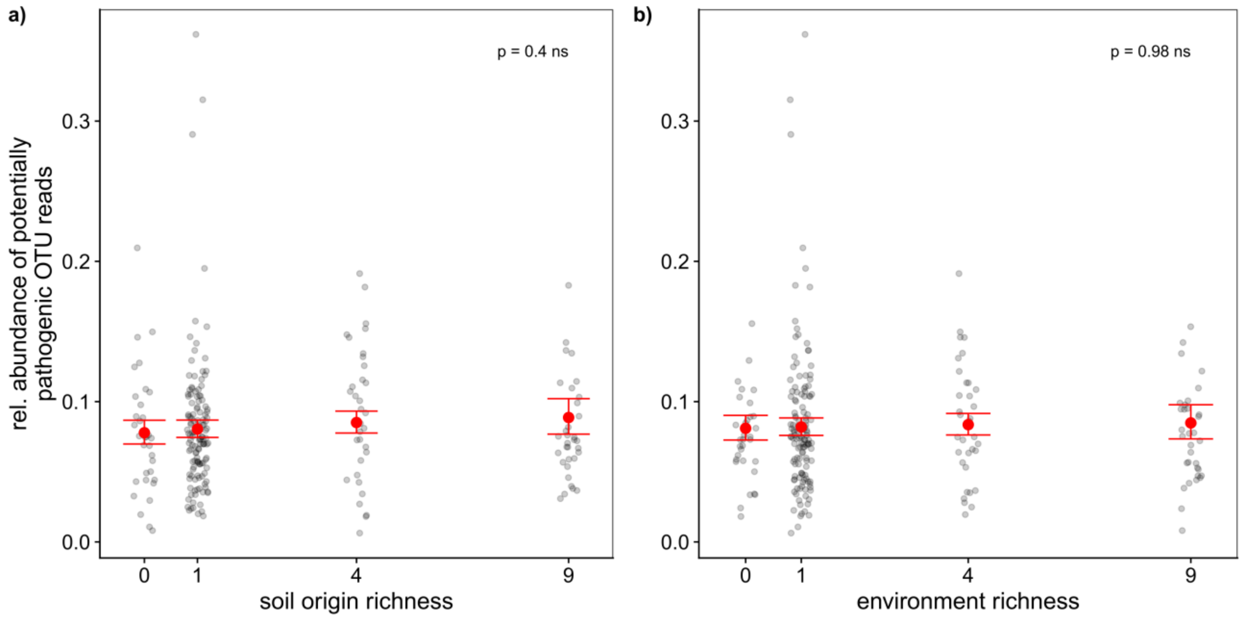


Figure S9: The relative abundance of potentially pathogenic OTU reads in the soil transplant experiment in response to a) soil-origin richness and b) environment richness. Raw data are in black and model prediction on the generalized linear mixed model (model 13) with the 95% interval are in red.

| Variable | Chi2 | P.Value | |
|---|---|---|---|
| log environment richness | 0.16 | 0.69 | |
| log soil origin richness | 46.10 | < 0.001 | *** |
| log e. rich. x log s.o. rich. | 2.44 | 0.12 | |

Table S21: Anova table to test for the overall effect of explanatory variables table for the generalized linear mixed-effects model of the probability for potentially symbiotic AMF OTUs to be present with log-transformed soil-origin richness, log-transformed environment richness and their interaction as fixed effects and plot identity as random effect (model 14).

| Fixed.Effects | Estimate | SE | z-value | P.value |
|---|---|---|---|---|
| intercept | -1.516 | 1.314 | -1.15 | 0.25 |
| log environment richness | -1.293 | 0.997 | -1.30 | 0.19 |
| log soil origin richness | 0.618 | 0.935 | 0.66 | 0.51 |
| log e. rich. x log s.o. richness | 1.079 | 0.729 | 1.48 | 0.14 |
| Random effects | Variance | SD | | |
| plot | 0.0217 | 0.147 | | |

Table S22: Summary table for the generalized linear mixed effects model of the probability for potentially symbiotic AMF OTUs to be present with log-transformed soil-origin richness, log-transformed environment richness and their interaction as fixed effects and plot identity as random effect (model 14).

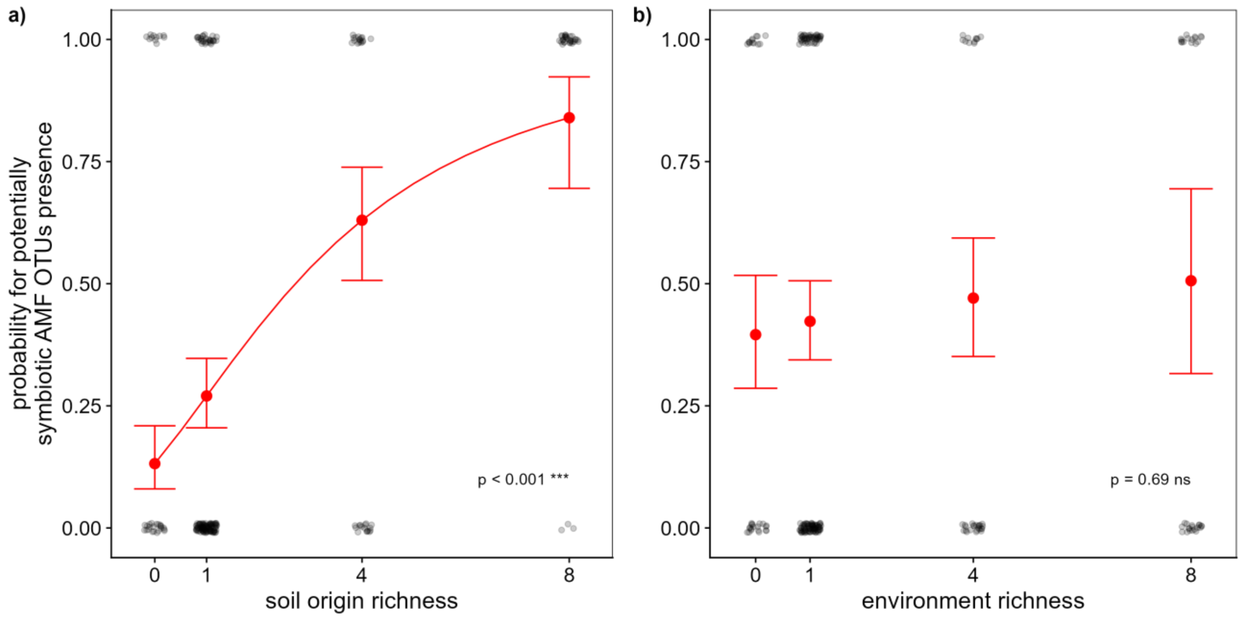


Figure S10: The probability for potentially symbiotic AMF OTUs to be present in the soil transplant experiment in response to a) soil-origin richness and b) environment richness. Raw data are in black and model prediction on the generalized linear mixed effects model (model 14) with the 95% interval are in red. Significant effects are shown as solid lines and for non-significant effects the model predictions are shown without a line connecting them.
